# *Toxoplasma* strikes preemptively to swiftly suppress macrophage immune response during active infection

**DOI:** 10.1101/2025.01.31.635863

**Authors:** Dominykas Murza, Filip Lastovka, George Wood, Matthew P. Brember, Ollie Chan, James W. Ajioka, Betty Y. W. Chung

## Abstract

The apicomplexan parasite *Toxoplasma gondii* is known to manipulate its host in multiple ways, ranging from proteins secreted into the host cell to hormone balance disruption and behavioral changes. Host immune system is crucial in managing the outcome with macrophages as part of the first line of defense. However, the initial triggers that ultimately result in characteristic complex responses and, at times, health hazards, remain poorly understood. This study focuses on filling the gaps in our knowledge of acute transcriptomic changes taking place in a mouse macrophage *T. gondii* infection model. We performed time-resolved transcriptomic profiling to simultaneously capture host and parasite gene expression profiles during the course of infection, focusing on the initial time frame of fifteen to 120 minutes, a crucial window for the host to activate innate immunity but also for the parasite to establish within the host macrophage. Further, utilization of inactivated parasite stimulation enabled dissection of transcriptomic response to active parasite infection from innate immune responses. Here, we observed that macrophages upregulate transcripts encoding suppressors of cytokine signaling by 30 minutes, specific to live parasite infection. Additionally, both pro-growth and stress marker genes were dysregulated. Concurrently, transcriptional response of *T. gondii* was milder in magnitude, with initial changes pointing at increasing transcription and growth capacity, followed by a delayed transcriptional response pertaining to secreted proteins. Taken together, these results demonstrate that macrophages mount a rapid transcriptional response upon active invasion by *T. gondii*. In contrast, the delayed transcriptional activation in the invading *Toxoplasma* highlights its reliance on alternative regulatory mechanisms to establish its replicative niche within the host.

**Author summary:** *Toxoplasma gondii,* a eukaryotic intracellular parasite, is often regarded as one of the most globally successful parasites because it can infect virtually any warm-blooded animal. It uses a repertoire of secretory proteins to gain a foothold in a host cell, often resulting in a dormant infection *in vivo* due to sufficient immune suppression. However, the timing of the signaling events as *Toxoplasma* invades is not yet fully understood. In this work we implement our user-friendly transcriptomic method to simultaneously capture *T. gondii* and murine macrophage protein-coding RNA contents over a time course to track cellular responses during infection. In particular, we focus on the first 2 hours of infection, a time window where the initial transcriptomic changes within macrophage are generally expected to take place and potentially define further course of infection. Our analysis reveals a robust host macrophage immune response and a moderate more gradual *T. gondii* response. These findings complement the currently existing picture of all the cellular regulation layers involved in *Toxoplasma*–macrophage interaction.

## Introduction

*Toxoplasma gondii* is a protozoan intracellular parasite that infects warm-blooded animals. Its importance is illustrated by its very high prevalence, with estimated 25.7% of global human population being seropositive[1]. In immunocompromised individuals, frequent sites of harm by *T. gondii* are immune-privileged organs, such as eye or brain[2]. Between the first exposure and the resulting tissue damage, many layers of regulation of cellular function, studied to different extent, act simultaneously, adding up to difficult-to-predict infection outcomes. We hypothesize that transcriptome disturbance during the infection event and immediately afterwards plays a substantial role in determining its outcome.

Previous studies have addressed multiple aspects of transcriptomics after infection by *T. gondii.* While *Toxoplasma* is known to infect a diverse repertoire of cell types, there are indications that transcriptional response to infection depends on the host cell type[3]. For example, in Vero cells, several infection-induced apoptosis pathways were activated[4]. Infection of human foreskin fibroblasts (HFF) by *T. gondii* bradyzoites resulted in upregulation of c-Myc, NF-κB and energy-related pathways[5]. *In vivo,* transcriptomic analyses of *T. gondii-*infected mouse brain tissue showed that infection disrupts normal expression patterns of genes associated with neurological functions[6], and that this observation also held true *in vitro*[7]. Another *in vivo* mouse brain study also demonstrated immune-related gene upregulation at both 11 and 33 days post infection with *T. gondii* oocysts[8]. Similar studies examined a variety of other organs, including testes and uterus[9], lung, lymph nodes and spleen[10], brain[11], and liver[12].

However, abundant lines of evidence point at the importance of different cell types, especially macrophages during infection by *T. gondii*[13]. The lineup of microneme (MIC), dense granule, and rhoptry proteins (ROP) secreted into the host upon infection has received particular attention. For example, the protein MIC3 caused increased TNF-α production and M1 macrophage activation[14]. The dense granule protein TgIST reaches the host cell nucleus to repress STAT1-dependent promoters[15]. ROP5, ROP17 and ROP18 collectively inhibit Immunity-Related GTPase (IRG) recruitment to the parasitophorous vacuole (PV)[16]. ROP16 can induce activation of STAT3 and STAT6, thereby initiating M2 polarization, a program with reduced anti-microbial capacity, to which suppressed generation of reactive oxygen species (ROS) is known to contribute[17]. Of note, macrophages can be subdued even without being infected, as secretory protein injection can occur regardless[18]. Finally, macrophages and their precursors, monocytes, also function as vehicles for facilitated migration of the parasite within the host’s body[19,20], which further highlights the importance of studying macrophage gene expression changes during *T. gondii* infection. The scarce transcriptomic evidence of immune cell interactions with *T. gondii* showed that the type I strain RH perturbed the transcription of mouse bone marrow-derived macrophages (BMDMs) more markedly than the type II strain PTG[21]. Yet, macrophage transcriptomics upon *T. gondii* infection has not been previously covered in greater detail.

In the intricate interplay between host and pathogen during entry, understanding the temporal dynamics of gene expression is crucial. Distinct alterations to both host and *Toxoplasma* transcriptomes were observed in various time scales following infection. In particular, expression patterns were previously analyzed from two hours to 32 days post-infection and beyond[3–7,10–12,21–23]. Notably, most studies to date have concentrated on the pre-infection conditions and the experimental endpoints, leaving the dynamic events occurring in between largely unexplored. Significant changes in the host transcriptome are anticipated within the first two hours post-infection, as demonstrated by studies involving viral[24], bacterial[25], and eukaryotic pathogens[26]. However, no comparable early-infection transcriptomic time-course data have been published for *T. gondii* or any host cell type it infects. We expect that during this critical period of initial interaction, the transcriptomes of *Toxoplasma* and macrophage undergo primary changes that propagate the response via effector and regulator molecules and ultimately contribute to determining the parasite’s establishment within the host. However, studying related phenomena *in vivo* is not feasible due to sparse infection and imprecise timing control. Our *in vitro* study design addresses asynchronous infection and dilution of signal caused by low infection rates. The obtained results highlight the importance of global transcriptional remodeling by *T. gondii* infection, particularly on early onset of signaling and transcription factors. We speculate that the observed extensive immediate transcriptional response directs further course of macrophage infection by *T. gondii*.

## Results

### Establishment of efficient *in vitro* infection

A caveat of many past experimental designs is the unknown degree of confounding influence on the transcriptome caused by non-specific activation (e.g. due to temperature changes or extracellular factors). It is known that macrophage transcriptome undergoes drastic changes in early-response cytokine gene expression as soon as one hour post-lipopolysaccharide (LPS) stimulation[27]. Likewise, *in vitro* infections are prone to include the non-specific stimulation component caused by lysed *T. gondii* in the medium. Extracellular single-stranded RNA, for instance, is expected to be present in such lysates and is known to activate Toll-like receptors (TLR) 7 and 8 in mice and humans, respectively[28]. Additionally, in the human myelomonocytic cell line THP-1, *T. gondii* lysate was also able to trigger immunosuppression via TLR2-mediated NF-κB activation[29]. Our design implements fully inactivated *T. gondii* lysate controls (further referred to as Dead *Toxoplasma*; Fig. S1) to estimate the extent of and adjust for the side effects of non-specific stimulation during *in vitro* infections as well as media-only controls (Mock) to account for any possible signal due to temperature fluctuations, mechanical stress etc.

The experimental workflow is outlined in Fig. 1A. For the procedure, we selected a common model mouse macrophage cell line RAW264.7 to facilitate comparison to previous studies[14,30,31] as well as Type I GFP-tagged *T. gondii* strain RH. We aimed to achieve adequate infection synchronicity as a prerequisite for this workflow. Therefore, the duration of parasite suspension exposure to RAW264.7 was set to 15 min to match the time scale of intended sampling resolution. Parasite concentration in a fixed volume of medium during infection was then optimized to yield at least 50% infection efficiency (Fig. 1B). Then, the selected concentration of 6×10^7^ tachyzoites/mL was used for the infection time course (15, 30, 60, 120 min and 24 h). For each time point, RAW264.7 cells were treated with either *Toxoplasma*-free medium (Mock), live *T. gondii* (Live *Toxoplasma*) and parallel dead *T. gondii* through successive freeze thaw cycles (Dead *Toxoplasma*) in triplicate. For the Live *Toxoplasma* condition, threshold infection efficiency was achieved at every time point (Fig. 1C). At 24 hours, increased burden of parasites leading to drastic changes in RAW264.7 morphology was observed. In parallel to imaging, sample total RNA was harvested, its quality was confirmed (Fig. S2), and two replicates were used to prepare PARFA-seq libraries.

**Figure 1.**
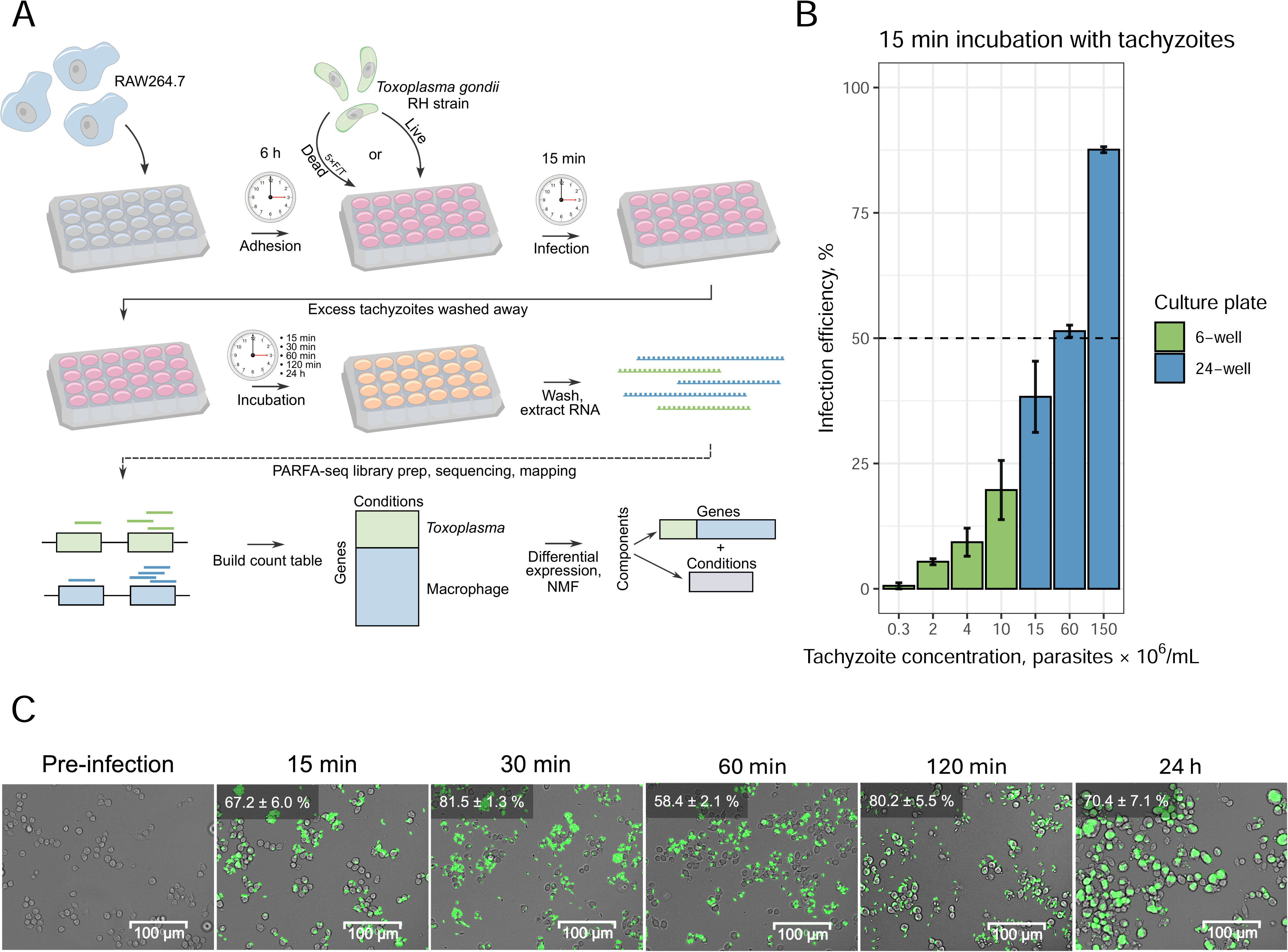
**A.** Experimental scheme of infection time course and downstream workflow; **B.** Optimization of tachyzoite concentration to achieve adequate infection rates, with a threshold set at 50% infection efficiency. Error bars represent ± 1 standard deviation; **C.** Infection time course micrograph merged from brightfield and green channels. *T. gondii* exhibits green fluorescence due to stable GFP expression in the strain used. Percentage at the top left indicates infection efficiency (± 1 standard deviation) for each time point.

### Polyadenylated RNA Fragment Abundance Sequencing library preparation and composition analysis

PolyAdenylated RNA Fragment Abundance (PARFA)-Seq was established to facilitate our understanding of transcriptome dynamics during *T. gondii* infection of macrophages. PARFA-seq library preparation workflow is illustrated in Fig. 2A. This open-source library preparation protocol is time-efficient, features dual indexing for multiplexing many samples and was tested to perform well with reagents commonly available in molecular biology laboratories (Fig. S3). Briefly, it involves reverse transcription of poly-A(+) RNAs, second strand synthesis by random priming and two rounds of PCR to attach necessary handles. Library quality control is also simple to assess through standard non-denaturing gel electrophoresis (Fig. S4). As *T. gondii* transcriptome is also polyadenylated, both the host and the parasite transcriptomes were simultaneously captured. Upon demultiplexing and adapter trimming, reads are mapped to a combined *T. gondii–*mouse genome and unique mappers are used for count matrix generation.

**Figure 2.**
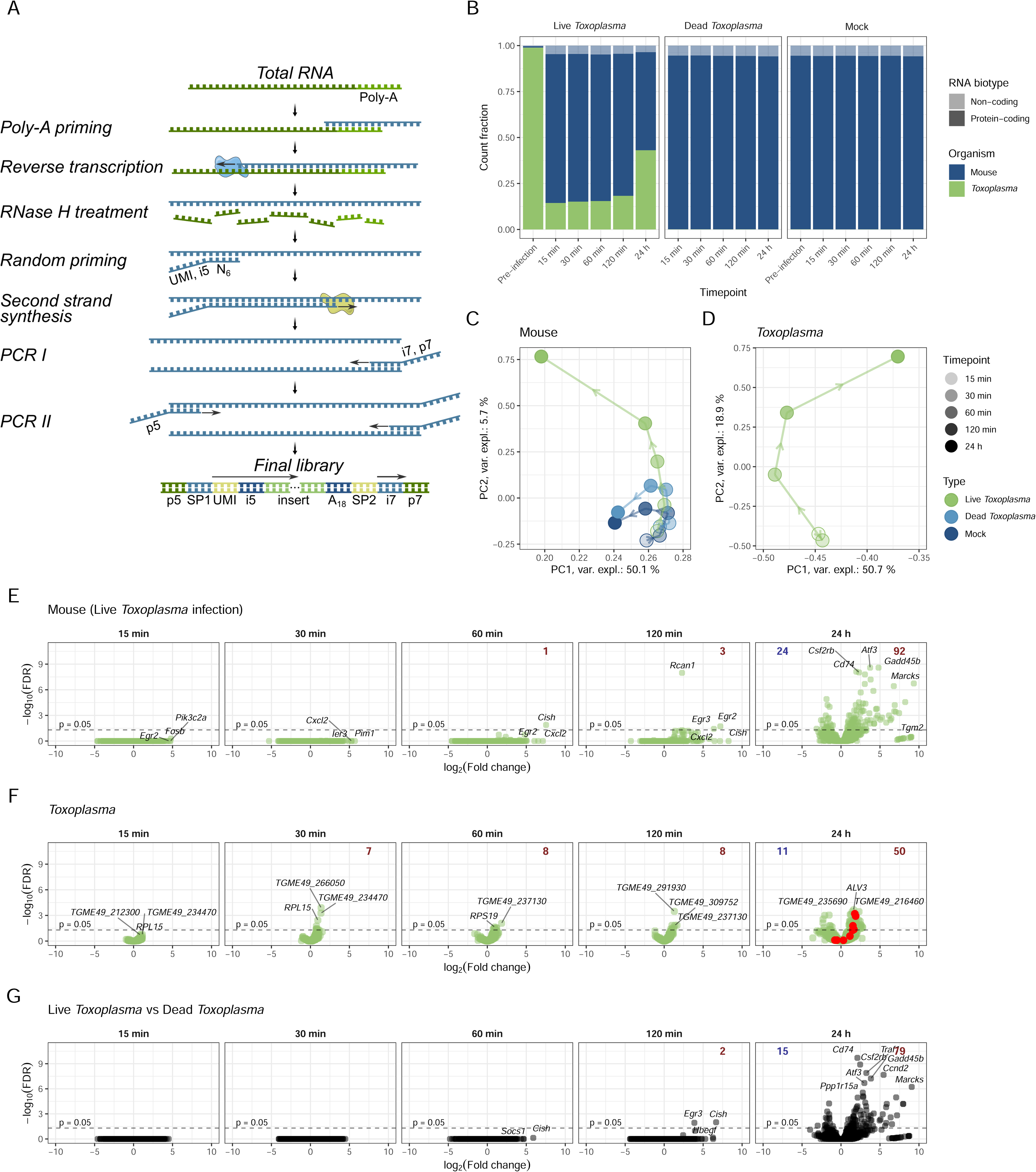
**A.** Library preparation strategy: poly-A(+) RNAs are captured by complementarity to reverse transcription primer, then second strand is synthesized using random hexamers and handles for sequencing are attached through two rounds of PCR. **B.** Mapped read distribution by species and by biotype. **C** and **D.** Principal component analysis of mouse and *T. gondii* transcriptomes, respectively. **E** and **F.** Volcano plots of Live *Toxoplasma*-infected mouse and *T. gondii* time points, respectively, compared to corresponding samples (either macrophage or *Toxoplasma*) before infection. Most significant and most upregulated transcripts are labelled. Number of significantly (p ≤ 0.05) downregulated and upregulated genes for each time point is given in blue and red, respectively. In F, 24-hour time point, red dots mark ROP genes. **G.** Volcano plot for macrophage transcriptomes when Live *Toxoplasma* and Dead *Toxoplasma* are compared for each timepoint.

Mapped read classification by species and RNA biotype (Fig. 2B) demonstrates species- and biotype-specific alignment. Additionally, increasing fraction of *T. gondii* transcript counts throughout the time course confirmed proliferation of the parasite and agrees with the micrographs (Fig. 1C) and the total RNA profiles (Fig. S2). Also, macrophages incubated with Dead *Toxoplasma* did not contain *T. gondii* RNA, confirming successful inactivation of *T. gondii*. Global transcription assessment through principal component analysis (PCA) revealed Live *Toxoplasma*-infected macrophage transcriptome notably diverged from both control conditions by 60 min (Fig. 2C). In parallel, significant change of *T. gondii* transcriptome at 60 min was also observed, whereas global difference between 15 min and 30 min less pronounced (Fig. 2D). Differential expression analysis was conducted by comparing each time point to the pre-infection state (Fig. 2E and F). Notably, transcriptional upregulation was more prominent than RNA decay for both the host and the parasite throughout the course of infection, as shown by higher numbers of significantly upregulated genes. In macrophages, typical early response genes that modulate inflammation (*Egr2, Fosb, Cxcl2, Cish*) were induced the most. In addition, comparing macrophage transcriptomes between Live and Dead *Toxoplasma* conditions enabled precise adjustment for non-specific stimulation, revealing that the suppressor of cytokine signaling response (*Cish*, *Socs1*) is specific to Live *Toxoplasma* infection (Fig. 2G). Among the highly upregulated *T. gondii* genes, no clear patterns or commonalities emerged throughout the time course. Statistically significant changes in *Toxoplasma* transcriptome were modest between 15–60 min post infection indicating similarity to the control condition - pre-infection *Toxoplasma* (Fig. 2F).

### *Toxoplasma* transcriptional landscape during macrophage infection

The behavior of eight groups of *T. gondii* genes known to be important for infection and parasite proliferation was initially examined throughout the time course. Fold changes at each time point were calculated relative to the pre-infection condition (time zero) to account for baseline expression and allow direct comparisons across all conditions and time points (Fig. 3A). Among the most populous gene groups shown, *T. gondii* underwent an increase in RNA polymerase gene expression at 60 min. Rhoptry and rhoptry neck (RON) proteins were upregulated at 24 hours (Fig. 2F, Fig. 3A).

**Figure 3.**
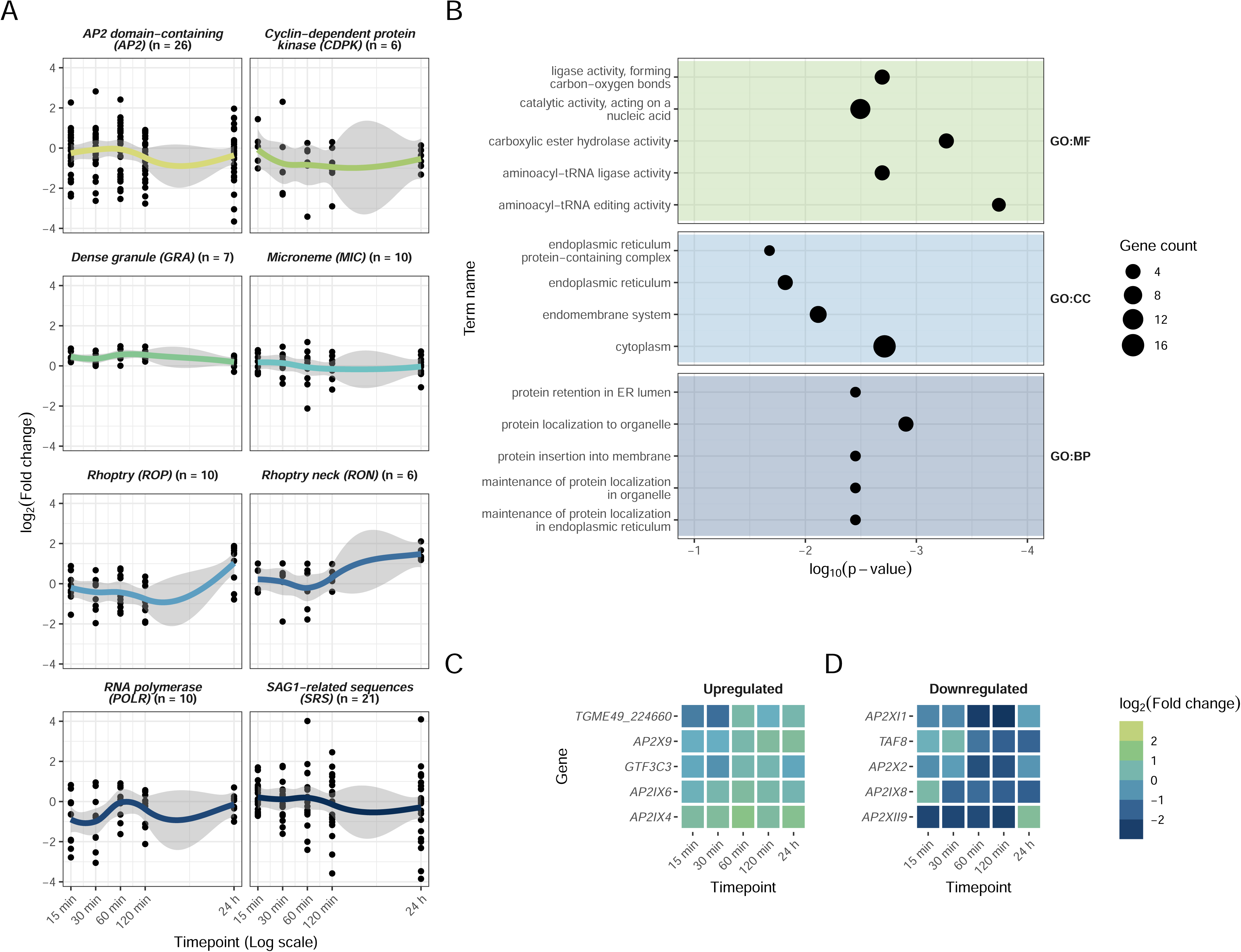
**A.** *T. gondii* gene group changes throughout the time course, expressed as log_2_ fold change relative to pre-infection tachyzoites. Grey area depicts the 95% confidence interval. **B.** Gene set enrichment plot for most enriched terms from *T. gondii* transcriptome features at 15 min to 120 min. Five most significant terms are displayed for each database. **C** and **D.** Transcriptional profiles of top five upregulated and downregulated *T. gondii* transcription factors, respectively.

An independent Non-negative Matrix Factorization (NMF) analysis was additionally performed on macrophage-*T. gondii* metatranscriptome from 15 to 120 minutes post-infection (excluding the 24-hour timepoint) to disentangle early responses (Fig. S6). Genes associated with transfer RNA charging function, cytoplasm localization and membrane-associated proteins were upregulated at early time points (Fig. 3B). Additionally, the differential expression of *T. gondii* transcription factors appeared relatively modest, with log_2_(fold change) of all genes falling within -3 to 3 (Fig. 3C and D). Collectively, these findings suggest an increased capacity for growth and reproduction within the host, as well as preparation for subsequent infection.

### Macrophage transcriptome response

We next focused on the macrophage response. In agreement with principal component analysis, NMF analysis showed subtle differences between Dead *Toxoplasma*-stimulated macrophages and mock-treated cells, whereas Live *Toxoplasma* condition was clearly distinct in component 3 (Fig. 4A). Gene set enrichment analysis of the driving features in each component (Fig. S7, Fig. 4B) revealed that functional enrichment – including responses to stimuli, transcription regulation, and immune processes – is concentrated in component 3.

**Figure 4.**
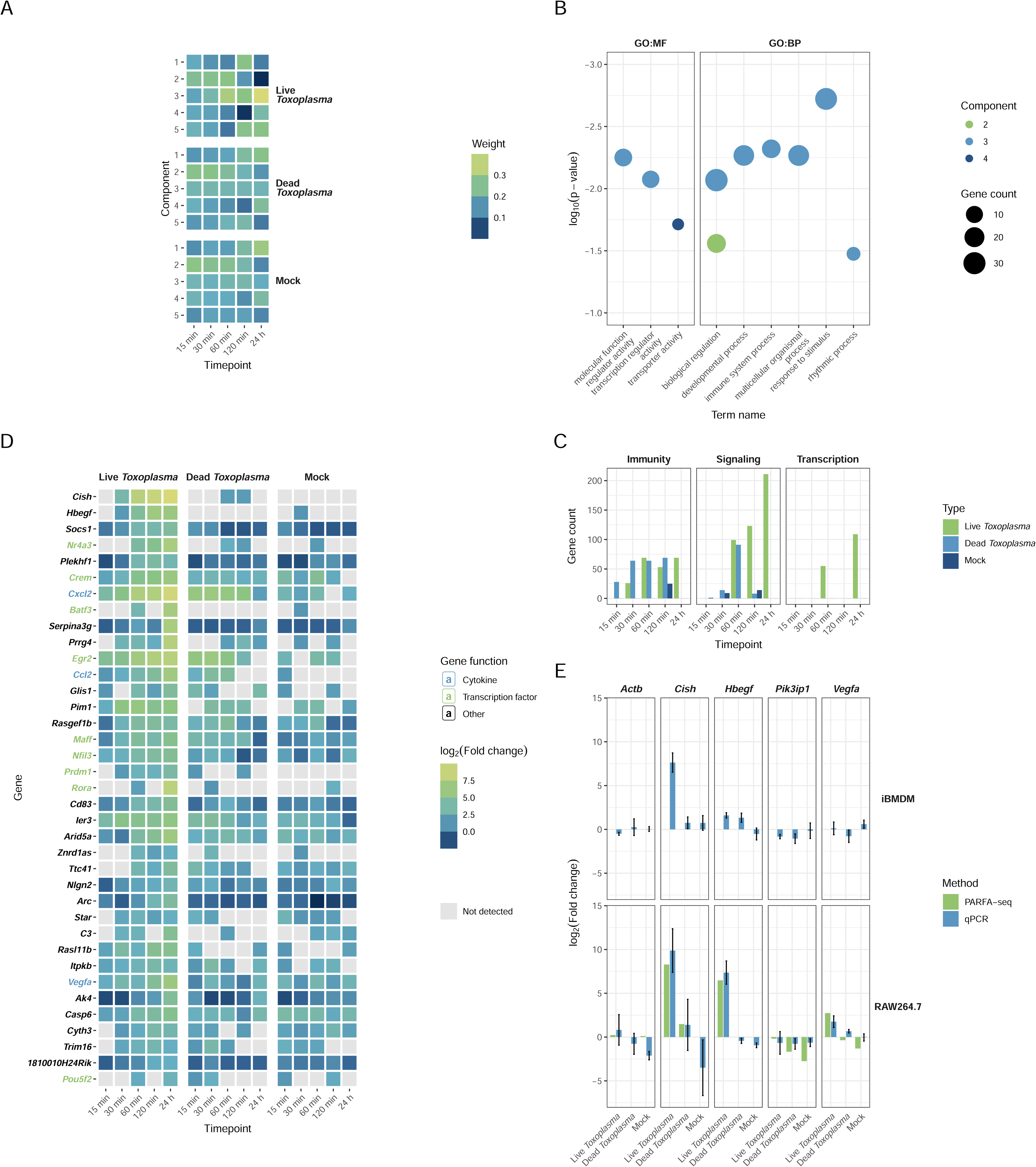
**A.** Mouse transcriptome non-negative matrix factorization component weights across all time points and conditions. Larger weight values indicate components with upregulated genes dominating. **B.** Gene set enrichment plot for selected features from each component in panel A. Components 1 and 5 did not exhibit enrichment in these broad terms. **C.** Number of genes in enriched gene ontology terms related to immunity, signaling, and transcription for each time point and condition. **D.** Time course transcriptional profiles for mouse genes characteristic to Component 3. **E.** PARFA-seq validation by quantitative PCR and gene expression changes during immortalized bone marrow-derived macrophage infection by *T. gondii* assessed by quantitative PCR.

To investigate the dynamics of immunity-, signaling-, and transcription-related genes, we focused on genes with a log_2_ fold change above two in each condition and counted those belonging to the enriched terms (Fig. 4C and Table S2). Transcription- and signaling-related gene enrichment was most pronounced in Live *Toxoplasma*- treated macrophages. However, during first two hours, Dead *Toxoplasma* also induced a comparable range of immune genes, in some cases preceding the response in Live *Toxoplasma-*infected macrophages.

Features characteristic of component 3 include suppressors of cytokine signaling, *Cish* and *Socs1*, both of which underwent marked upregulation, specific to Live *Toxoplasma* infection (relative to Dead *Toxoplasma* treatment) by 30 minutes (Fig. 4D). This observation corroborates previous evidence that *Cish* and *Socs1* are upregulated at four hours post-infection[32]. Interestingly, both Live and Dead *T. gondii* treatments trigger cytokine (*Cxcl2*, *Ccl2*) upregulation, although this response was delayed during live *Toxoplasma* infection. Among the main transcription factors driving the early (<60 min) transcriptional response is *Nr4a3*, in line with previous findings[33].

Finally, expression pattern of a range of genes, including *Cish,* identified through PARFA-seq were validated by qPCR, underscoring the robustness of PARFA-seq in detecting transcriptional changes (Fig. 4E). The same targets were also assessed by qPCR in an equivalent murine immortalized bone marrow-derived macrophage (iBMDM) infection time course, which largely recapitulated the characteristic macrophage response to *T. gondii* infection.

## Discussion

In this work, we applied PARFA-seq to investigate the transcriptomic responses of mouse macrophages and *T. gondii* immediately following invasion. Our results show that macrophages undergo a pronounced transcriptional response within the first 30–60 minutes of parasite contact. These early changes are likely to be crucial in shaping the subsequent course of infection by suppressing host innate immunity to aid in establishment within the macrophage.

We have identified a set of genes involved in immune response, signaling and transcription regulation that act early in the infection timeframe. In line with previous findings [32], suppressors of cytokine signaling – *Cish* and *Socs1 –* were upregulated. Their well-established function is the provide negative feedback and prevent overactivation of the immune system[34]. However, they are also apparent targets for exploitation by various pathogens including Avian leucosis virus[35], Human immunodeficiency virus[36] or *Listeria monocytogenes*[37]. In under 30 min post-infection by the *T. gondii* RH strain, expression of another gene of the group, *Socs3,* was also observed at the protein level[38], likely reflecting manipulation by the parasite rhoptry kinase ROP16 that phosphorylates STAT3, leading to its translocation to nucleus and pro-growth response[39]. The more pronounced upregulation of *Socs1* relative to *Socs3* in our dataset implies that M2 macrophage polarization due to *T. gondii* infection is favoured[40].

Additionally, our data revealed upregulation of other genes linked to proliferation. Notably, *Hbegf* is known to bind epidermal growth factor receptors (EGFR) to promote cell cycle progression and survival[41] and *Vegfa* facilitates macrophage migration[42]. Recently, *Nr4a3* was also labelled as an important mediator of increased migration through induction by ROP28[33]. Taken together, these responses may help create conditions conducive to parasite survival, while simultaneously protecting the host cell from early cell death, potentially aiding parasite dissemination. Nevertheless, genes related to anti-infection processes such as autophagy regulation (*Plekhf1*) or neutrophil recruitment (*Cxcl2*)[43] were also induced. Despite these defenses, infection ultimately becomes established, indicating that the macrophage’s early response is not sufficient to halt *T. gondii.* In our dataset, these effects appear specific to live *T. gondii* infection, suggesting that the parasite triggers them through secreted factors and intracellular mechanisms. Other rapidly acting layers of regulation – such as translational control[44] – may also contribute, but remain to be explored.

Meanwhile, the transcriptional response of the invading *T. gondii* tachyzoite itself appears to be relatively modest and reflects a gradual shift in increased growth-related capabilities. We did not observe sudden decrease in *T. gondii* transcripts, nor surge of transcription, presumably because many secreted effector proteins required for immune evasion are already synthesized prior to invasion. Thus, the parasite effectively “pre-arms” for infection, and the outcome may be largely predetermined by these secreted factors prior host-parasite interaction. As a result, rhoptry-related genes (ROP and RON) are upregulated more slowly, as evidenced by our 24-hour time point. Overall, the parasite appears to rely on a steady, business-as-usual approach, rather than undergo a rapid, finely orchestrated, and environment-tailored shift of transcriptional program.

Several avenues warrant further investigation and harnessing single-cell transcriptomics represents a promising way to address the current limitations in our approach, particularly in light of variable parasite infectivity. Although our design maximizes the fraction of invaded macrophages, it also increases the number of *T. gondii* tachyzoites that fail to infect, potentially diluting the parasite-specific transcriptomic signal. Future efforts specifically dedicated to the parasite’s response might therefore consider lower parasite-to-host ratios (bulk analyses) or omit uninfecting parasite cells (single-cell resolution). Additionally, while *in vitro* assays offer targeted mechanistic insights, they inevitably overlook the complexity of *in vivo* infections that involve multiple cell types and genetic variability in both host and parasite.

By comparing live *T. gondii* infection with dead-parasite stimulation, our study provides a unique perspective on the rapid host response. We show that macrophages mount a pronounced immune reaction by the first hour, but this response is dampened specifically in the presence of live parasites, facilitating *T. gondii* establishment within the macrophage. This insight underscores the importance of early host-parasite interactions in shaping the outcome of infection and suggests new directions for dissecting these events.

## Methods

### Cell cultures

GFP-tagged *Toxoplasma gondii* type I strain RH (acquired from Frickel Lab) was used for all infection experiments. Human foreskin fibroblast (HFF) cells (ATCC) were used for routine *T. gondii* maintenance and expansion for infections. Mouse macrophage-like cell line RAW264.7 (ATCC) was passaged by scraping off, resuspension and 10-fold dilution when the culture reached approximately 80% confluence, which was typically every 3 days. HFFs were grown to near 100% confluence and passaged after trypsinization and 5- to 10-fold dilution. 100% confluence after passaging is achieved in approximately 1 week. Fully confluent HFFs were maintained for up to 2 weeks for further use. *Toxoplasma gondii* was maintained in HFFs at 90-100% confluence by passaging 10–100 µL (based on approximate visual estimation of parasite counts) of media containing extracellular tachyzoites from an infected T25 flask to an uninfected T25 flask. This procedure was carried out every 2–4 days. All uninfected and infected mammalian cell cultures were grown in Dulbecco’s Modified Eagle’s Medium – High Glucose (DMEM-HG) (Sigma) with addition of 10% initial volume of Fetal Bovine Serum (FBS; Gibco™) and 1% initial volume of 200 mM L-glutamine (Gibco™) in a 37 °C incubator with a 5% CO_2_ atmosphere and 100% relative humidity. All mammalian cultures were routinely restarted from new cryostocks after they have reached passage number 15. All procedures involving work with mammalian cell lines and *T. gondii* were carried out in a BSL2 facility.

### *T. gondii* preparation for infection time course

Before an infection experiment, *T. gondii* tachyzoite cultures in HFFs were expanded sequentially. First, when approximately 80% of HFFs have been lysed by the parasites in the T25 maintenance flask, a small aliquot was taken for further passaging while all the remaining extracellular parasites were used to infect a T75 flask of confluent HFFs. After 80% of HFF cells have lysed (typically between 24 and 48 hours post-infection), the culture supernatant was distributed between three T175 flasks with confluent HFFs. Finally, after another 24–48 hours, approximately 20% of HFF monolayer was lysed, at which point intracellular *T. gondii* tachyzoites were harvested for infection. The culture supernatant is removed and the tachyzoite-laden HFF monolayer was scraped off, resuspended in 5 mL of supplemented DMEM (pre-warmed to 37 °C) for each T175 flask and transferred to a 15 mL polypropylene tube. The suspension was then passed through a 27G needle 4–6 times using gentle pressure on the plunger to break tachyzoites free. The parasites were then separated from debris by differential centrifugation: the suspension was centrifuged at 100 g for 5 min at room temperature, the supernatant was transferred to another tube followed by another centrifugation at 1000 g for 5 min at room temperature. The supernatant was carefully removed and the pellet containing the parasites was gently resuspended in 500 µL of pre-warmed medium. Parasite concentration was quantified by hemocytometry of a 100-fold dilution in phosphate-buffered saline (Gibco). One T175 HFF flask routinely yielded 2×10^8^ to 4×10^8^ purified intracellular tachyzoites. Dead *Toxoplasma* for control infection was prepared by performing five freeze-thaw cycles in dry ice–ethanol slurry and warm water. Full lysis of the parasites was confirmed by observation under a microscope.

### Macrophage infection

At approximately 80% confluence, RAW264.7 or iBMDM cells were mechanically scraped off the bottom of the flask and resuspended in 10 mL of pre-warmed medium and quantified using hemocytometry. A T75 flask routinely yielded about 4×10^7^ cells. Cells were diluted to the desired concentration with medium and seeded into multi-well tissue culture plates. 2.5 mL or 0.5 mL of medium was used for standard 6-well plates and 24-well plates, respectively. After seeding, the cells were allowed to adhere to the bottom of the plate for six hours to overnight before infection. One plate per time point was prepared. For infection, the medium was removed, and the parasite suspension (1.2×10^7^ tachyzoites in 200 µL of medium per well in a 24-well plate) was immediately added to the wells. For Mock or Dead *Toxoplasma* infections, 200 µL of medium or lysed *T. gondii,* respectively, were used. Then, the plates were stacked and gently rocked by hand in a forward-backward and then left-right motion for 30 s to distribute the added tachyzoites. The plates were placed in the incubator unstacked. After 15 min, the plates were removed from the incubator. This was considered time point zero for the infection[45]. The parasite suspension was removed from the wells followed by two washes (500 µL of pre-warmed medium each) to remove unattached tachyzoites. Another 500 µL of medium was added and further incubation was carried out in the incubator until the desired time point, when the medium was removed and a single wash with 500 µL of PBS was performed. After the removal of PBS, the wells were promptly imaged with ZOE Fluorescent Cell Imager (BioRad) in brightfield and green channels and the cells were immediately lysed by addition of 200 µL of TRIzol™ Reagent (Invitrogen™). The lysates were then stored at -20 °C. Total RNA was extracted according to manufacturer’s guidelines, dissolved in 10 mM Tris-HCl (pH 7.5) and quantified using NanoDrop™ One. RNA integrity was confirmed on 2% agarose TBE gels. The RNA was stored at -80 °C until further use.

### Infection efficiency quantification

To quantify the fraction of macrophages that were infected, micrographs from brightfield and green channels were merged with ImageJ and cells were manually counted using an ImageJ plugin Cell Counter[46]. Infection efficiency was defined as the following:

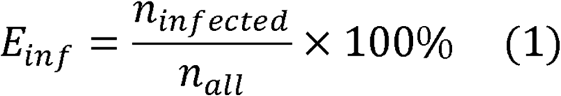

Where E_inf_ is the infection efficiency in percent, n_infected_ is the number of macrophages in a micrograph viewing field that colocalize with GFP fluorescence of the parasites, n_all_ is the number of all the macrophages present in that field. Micrographs were taken in the centre of each well. Three random non-overlapping fields were quantified, and the average infection efficiency value was calculated.

### qRT-PCR

Quantitative reverse transcription PCR was used to verify expected infection responses and validate RNA sequencing results. Total RNA was reverse-transcribed with M-MLV reverse transcription (RT) enzyme (Clontech), MMLV RT 5X buffer (Promega), and random hexamers (Promega) in the presence of RNaseOUT™ (Invitrogen™) ribonuclease inhibitor. The synthesized cDNA was diluted five times. 2 µL of cDNA per well were used in 384-well plates (Applied Biosystems), along with iTaq™ Universal SYBR® Green Supermix (BioRad) and 0.5 µM final concentrations of forward and reverse primer (for primer sequences see Table S1). Primers against *T. gondii* genes were designed using NCBI Primer-BLAST to span exon-exon junctions. Reference gene and target gene reactions for three biological replicates per condition were performed in Applied Biosystems ViiA™ 7 Real-Time PCR System on fast mode. Analysis was performed using the ΔΔCt method[47] comparing 15 min and 120 min timepoints. Assays were controlled for species specificity, genomic DNA amplification and ambient contamination.

### PARFA-Seq library preparation

Libraries for transcript quantification by PARFA-Seq were prepared using total RNA from the RAW264.7 infection time course. The primer 3PS-RTP was annealed to poly-A(+) RNA, which was then reverse transcribed with SuperScript™ III (Invitrogen™). Samples were then treated with RNase H (Thermo Scientific™). Unincorporated oligonucleotides were removed using Sera-Mag™ magnetic beads (Sigma) according to manufacturer’s protocol. Oligonucleotide PARFA-uRPI-# containing a random hexanucleotide at the 3′ end was annealed to the cDNA, allowing the synthesis of the second DNA strand using NEBNext® Ultra™ II Q5® Master Mix (NEB) followed by addition of RPI-# and PCR for five cycles. Each library was prepared with a unique combination of PARFA-uRPI-# and RPI-#. PCR products were then size-selected using Sera-Mag™ magnetic beads at 0.65 and 1.00 ratio of beads to sample by volume. DNA after size selection was then amplified using primers RP1 and RPI-# using the same master mix for another 12–15 cycles, followed by second round of size selection using the same parameters. Library quality was verified by TBE-PAGE in 0.75 mm 6% gels. Gel images were used to estimate DNA concentration based on band intensity in the region of interest (approximately 350 to 500 bp). Image Studio™ Lite software (LI-COR Biosciences) was used for this purpose. Equal amounts of each library were pooled (1 pool per replicate). Library pools were further size-selected by gel electrophoresis: 1 mm precast 10% TBE-PAGE gel (Invitrogen™) were loaded with pooled libraries, run for 40 min at 200 V. The gel was stained with SYBR™ Gold (Invitrogen™) for 5 min, visualized in blue LED light and 350–500 bp regions were excised and subjected to DNA extraction. Briefly, slices containing DNA of interest were crushed, extracted with 500 μL of DNA extraction solution (300 mM NaCl, 10 mM Tris pH 8.5, 1 mM EDTA) overnight. Next day, the supernatant was filtered off and mixed with 2 μL of GlycoBlue™ coprecipitant (Invitrogen™), 500 μL of isopropanol, and allowed to precipitate at -20 °C for 1 h to overnight. The DNA was pelleted by centrifugation (4 °C, >20 000 g for 1 h) and washed twice with cold 80% ethanol. The precipitate was then air-dried and dissolved in 10 μL of 10 mM Tris (pH 8.5, nuclease-free). The libraries were quantified with Qubit™ 4 fluorometer and dsDNA HS Assay Kit (Invitrogen™). Replicates 2 and 3 were pooled together in equimolar ratio and mixed with additional libraries to increase complexity. All library preparation procedures were performed in 0.5–2.0 mL DNA LoBind® tubes (Eppendorf), where applicable. A scheme of PARFA-Seq library preparation is given in Fig. 2A.

### Next-Generation Sequencing and read pre-processing

Libraries were sequenced in Illumina® NextSeq® 500 instrument (1 × 75 bp mode) by Cambridge Genomic Services. The reads were output from the Illumina® sequencing platform and grouped by the 3′ barcode. The resulting data were then sorted based on a 5′ barcode, which was then trimmed from the reads. All 5′ barcode trimming and UMI manipulation was done using *umi_tools*[48]. The remaining poly-A sequences (5 nt or longer) were removed with *bbduk*[49]. Filtered reads were aligned to the m10 genome of *Mus musculus*, combined with *Toxoplasma gondii* genome built with ME49 release 29 using *STAR* aligner[50]. PCR bias was accounted for using *umi_tools dedup*: if multiple reads with the same UMI are aligned in the same position, only one read is retained. To generate a read count table, each sample was passed through *htseq-count*[51] and reads that aligned to exactly one feature in the GTF file were counted.

### Downstream transcriptome analysis

Genes with fewer than ten counts over all samples were filtered out. *DESeq2* package[52] was used for differential expression analysis. To deconvolute expression data, non-negative Matrix Factorization (NMF) was selected because of intuitive interpretation as well as its ability to assign genes to every component quantitatively by their weight scores, as opposed to qualitative clustering (e.g. k-means, where each gene belongs to one cluster only). NMF R package[53] was used to cluster genes of similar dynamics into components and extract the dominant features in each component. Mouse and *T. gondii* gene sets of interest were screened for GO-term enrichment using *gprofiler2*[54] and *ToxoDB*[55], respectively.

## Data availability

Raw and processed data are available from ArrayExpress accession E-MTAB-14557. Customized scripts used for this project are available upon request.

## Acknowledgements

We would like to thank Eva Frickel (University of Birmingham) for providing *T. gondii* strains. Postgraduate studies of D.M. was funded by The State Studies Foundation of the Republic of Lithuania and Research Council of Lithuania. F.L. was supported by a BBSRC DTP studentship. G.W. was supported by the Department of Pathology PhD studentship. The B.Y.W.C. laboratory was supported by a Medical Research Council Fellowship [MR/R021821/1], BBSRC project grants [BB/X001261/1, BB/V017780/1 and BB/V006096/1] and a Royal Society Research Grant [RGS\R2\192222]. Assets by Servier and DBCLS from bioicons.com[56] were used for illustrations.

## Author contributions

B.Y.W.C. conceived the research. B.Y.W.C., and D.M. designed experiments. D.M. conducted experiments, generated PARFA-Seq libraries and performed most of the bioinformatic analysis. F.L. and O.C. conducted qPCR and iBMDM infections. J.A. revised the manuscript and provided *Toxoplasma* strains and protocols. M.B. sequenced libraries and performed data pre-processing and mapping. B.Y.W.C. and D.M. wrote the manuscript.

## References

1. A M, K N, M H, W W. Global status of Toxoplasma gondii infection: systematic review and prevalence snapshots. Trop Biomed. 2019;36. Available: http://mymedr.afpm.org.my/publications/97679

2. Fisch D, Clough B, Frickel E-M. Human immunity to Toxoplasma gondii. PLOS Pathog. 2019;15: e1008097. doi:10.1371/journal.ppat.1008097

3. Swierzy IJ, Händel U, Kaever A, Jarek M, Scharfe M, Schlüter D, et al. Divergent co-transcriptomes of different host cells infected with Toxoplasma gondii reveal cell type-specific host-parasite interactions. Sci Rep. 2017;7: 7229. doi:10.1038/s41598-017-07838-w

4. Du K, Lu F, Xie C, Ding H, Shen Y, Gao Y, et al. Toxoplasma gondii infection induces cell apoptosis via multiple pathways revealed by transcriptome analysis. J Zhejiang Univ-Sci B. 2022;23: 315–327. doi:10.1631/jzus.B2100877

5. Sugi T, Tomita T, Kidaka T, Kawai N, Hayashida K, Weiss LM, et al. Single Cell Transcriptomes of In Vitro Bradyzoite Infected Cells Reveals Toxoplasma gondii Stage Dependent Host Cell Alterations. Front Cell Infect Microbiol. 2022;12. Available: https://www.frontiersin.org/articles/10.3389/fcimb.2022.848693

6. Tanaka S, Nishimura M, Ihara F, Yamagishi J, Suzuki Y, Nishikawa Y. Transcriptome Analysis of Mouse Brain Infected with Toxoplasma gondii. Infect Immun. 2013;81: 3609–3619. doi:10.1128/IAI.00439-13

7. Mouveaux T, Roger E, Gueye A, Eysert F, Huot L, Grenier-Boley B, et al. Primary brain cell infection by Toxoplasma gondii reveals the extent and dynamics of parasite differentiation and its impact on neuron biology. Open Biol. 2021;11: 210053. doi:10.1098/rsob.210053

8. Hu R-S, He J-J, Elsheikha HM, Zou Y, Ehsan M, Ma Q-N, et al. Transcriptomic Profiling of Mouse Brain During Acute and Chronic Infections by Toxoplasma gondii Oocysts. Front Microbiol. 2020;11. Available: https://www.frontiersin.org/articles/10.3389/fmicb.2020.570903

9. Wang J, Liu T, Mahmmod YS, Yang Z, Tan J, Ren Z, et al. Transcriptome Analysis of Testes and Uterus: Reproductive Dysfunction Induced by Toxoplasma gondii in Mice. Microorganisms. 2020;8: 1136. doi:10.3390/microorganisms8081136

10. He J-J, Ma J, Wang J-L, Zhang F-K, Li J-X, Zhai B-T, et al. Global Transcriptome Profiling of Multiple Porcine Organs Reveals Toxoplasma gondii-Induced Transcriptional Landscapes. Front Immunol. 2019;10. Available: https://www.frontiersin.org/articles/10.3389/fimmu.2019.01531

11. Chen L-F, Han X-L, Li F-X, Yao Y-Y, Fang J-P, Liu X-J, et al. Comparative studies of Toxoplasma gondii transcriptomes: insights into stage conversion based on gene expression profiling and alternative splicing. Parasit Vectors. 2018;11: 402. doi:10.1186/s13071-018-2983-5

12. He J-J, Ma J, Elsheikha HM, Song H-Q, Huang S-Y, Zhu X-Q. Transcriptomic analysis of mouse liver reveals a potential hepato-enteric pathogenic mechanism in acute Toxoplasma gondii infection. Parasit Vectors. 2016;9: 427. doi:10.1186/s13071-016-1716-x

13. Park J, Hunter CA. The role of macrophages in protective and pathological responses to Toxoplasma gondii. Parasite Immunol. 2020;42: e12712. doi:10.1111/pim.12712

14. Qiu J, Xie Y, Shao C, Shao T, Qin M, Zhang R, et al. Toxoplasma gondii microneme protein MIC3 induces macrophage TNF-α production and Ly6C expression via TLR11/MyD88 pathway. PLoS Negl Trop Dis. 2023;17: e0011105. doi:10.1371/journal.pntd.0011105

15. Gay G, Braun L, Brenier-Pinchart M-P, Vollaire J, Josserand V, Bertini R-L, et al. Toxoplasma gondii TgIST co-opts host chromatin repressors dampening STAT1- dependent gene regulation and IFN-γ-mediated host defenses. J Exp Med. 2016;213: 1779–1798. doi:10.1084/jem.20160340

16. Etheridge RD, Alaganan A, Tang K, Lou HJ, Turk BE, Sibley LD. The Toxoplasma pseudokinase ROP5 forms complexes with ROP18 and ROP17 kinases that synergize to control acute virulence in mice. Cell Host Microbe. 2014;15: 537–550. doi:10.1016/j.chom.2014.04.002

17. Kochanowsky JA, Thomas KK, Koshy AA. ROP16-Mediated Activation of STAT6 Suppresses Host Cell Reactive Oxygen Species Production, Facilitating Type III Toxoplasma gondii Growth and Survival. mBio. 2021;12: 10.1128/mbio.03305-20. doi:10.1128/mbio.03305-20

18. Koshy AA, Dietrich HK, Christian DA, Melehani JH, Shastri AJ, Hunter CA, et al. Toxoplasma co-opts host cells it does not invade. PLoS Pathog. 2012;8: e1002825. doi:10.1371/journal.ppat.1002825

19. Courret N, Darche S, Sonigo P, Milon G, Buzoni-Gâtel D, Tardieux I. CD11c- and CD11b- expressing mouse leukocytes transport single Toxoplasma gondii tachyzoites to the brain. Blood. 2006;107: 309–316. doi:10.1182/blood-2005-02-0666

20. Da Gama LM, Ribeiro-Gomes FL, Guimarães U, Arnholdt ACV. Reduction in adhesiveness to extracellular matrix components, modulation of adhesion molecules and in vivo migration of murine macrophages infected with Toxoplasma gondii. Microbes Infect. 2004;6: 1287–1296. doi:10.1016/j.micinf.2004.07.008

21. Menard KL, Bu L, Denkers EY. Transcriptomics analysis of Toxoplasma gondii-infected mouse macrophages reveals coding and noncoding signatures in the presence and absence of MyD88. BMC Genomics. 2021;22: 130. doi:10.1186/s12864-021-07437-0

22. 22. Data Set Tachyzoite Transcriptome Time Series (RH). [cited 23 Nov 2023]. Available: https://toxodb.org/toxo/app/record/dataset/DS_6529b4f68d

23. Minot S, Melo MB, Li F, Lu D, Niedelman W, Levine SS, et al. Admixture and recombination among Toxoplasma gondii lineages explain global genome diversity. Proc Natl Acad Sci. 2012;109: 13458–13463. doi:10.1073/pnas.1117047109

24. Marcinowski L, Lidschreiber M, Windhager L, Rieder M, Bosse JB, Rädle B, et al. Real- time Transcriptional Profiling of Cellular and Viral Gene Expression during Lytic Cytomegalovirus Infection. PLOS Pathog. 2012;8: e1002908. doi:10.1371/journal.ppat.1002908

25. Aprianto R, Slager J, Holsappel S, Veening J-W. Time-resolved dual RNA-seq reveals extensive rewiring of lung epithelial and pneumococcal transcriptomes during early infection. Genome Biol. 2016;17: 198. doi:10.1186/s13059-016-1054-5

26. Rabhi I, Rabhi S, Ben-Othman R, Rasche A, Consortium S, Daskalaki A, et al. Transcriptomic Signature of Leishmania Infected Mice Macrophages: A Metabolic Point of View. PLoS Negl Trop Dis. 2012;6: e1763. doi:10.1371/journal.pntd.0001763

27. Das A, Yang C-S, Arifuzzaman S, Kim S, Kim SY, Jung KH, et al. High-Resolution Mapping and Dynamics of the Transcriptome, Transcription Factors, and Transcription Co-Factor Networks in Classically and Alternatively Activated Macrophages. Front Immunol. 2018;9. doi:10.3389/fimmu.2018.00022

28. Heil F, Hemmi H, Hochrein H, Ampenberger F, Kirschning C, Akira S, et al. Species- Specific Recognition of Single-Stranded RNA via Toll-like Receptor 7 and 8. Science. 2004;303: 1526–1529. doi:10.1126/science.1093620

29. Lee E-J, Heo Y-M, Choi J-H, Song H-O, Ryu J-S, Ahn M-H. Suppressed Production of Pro- inflammatory Cytokines by LPS-Activated Macrophages after Treatment with Toxoplasma gondii Lysate. Korean J Parasitol. 2008;46: 145–151. doi:10.3347/kjp.2008.46.3.145

30. Ma Z, Li Z, Jiang R, Li X, Yan K, Zhang N, et al. Virulence-related gene wx2 of Toxoplasma gondii regulated host immune response via classic pyroptosis pathway. Parasit Vectors. 2022;15: 454. doi:10.1186/s13071-022-05502-5

31. Dong K, Jiang Z, Zhang J, Qin H, Chen J, Chen Q. The role of SIRT1 in the process of Toxoplasma gondii infection of RAW 264.7 macrophages. Front Microbiol. 2022;13. doi:10.3389/fmicb.2022.1017696

32. Zimmermann S, Murray PJ, Heeg K, Dalpke AH. Induction of Suppressor of Cytokine Signaling-1 by Toxoplasma gondii Contributes to Immune Evasion in Macrophages by Blocking IFN-γ Signaling. J Immunol. 2006;176: 1840–1847. doi:10.4049/jimmunol.176.3.1840

33. Hoeve AL ten, Braun L, Rodriguez ME, Olivera GC, Bougdour A, Belmudes L, et al. The Toxoplasma effector GRA28 promotes parasite dissemination by inducing dendritic cell- like migratory properties in infected macrophages. Cell Host Microbe. 2022;30: 1570–1588.e7. doi:10.1016/j.chom.2022.10.001

34. Elliott J, Johnston JA. SOCS: role in inflammation, allergy and homeostasis. Trends Immunol. 2004;25: 434–440. doi:10.1016/j.it.2004.05.012

35. Feng M, Xie T, Li Y, Zhang N, Lu Q, Zhou Y, et al. A balanced game: chicken macrophage response to ALV-J infection. Vet Res. 2019;50: 20. doi:10.1186/s13567-019-0638-y

36. Ryo A, Tsurutani N, Ohba K, Kimura R, Komano J, Nishi M, et al. SOCS1 is an inducible host factor during HIV-1 infection and regulates the intracellular trafficking and stability of HIV-1 Gag. Proc Natl Acad Sci. 2008;105: 294–299. doi:10.1073/pnas.0704831105

37. Mishra R, Krishnamoorthy P, Kumar H. MicroRNA-30e-5p Regulates SOCS1 and SOCS3 During Bacterial Infection. Front Cell Infect Microbiol. 2021;10. doi:10.3389/fcimb.2020.604016

38. Whitmarsh RJ, Gray CM, Gregg B, Christian DA, May MJ, Murray PJ, et al. A Critical Role for SOCS3 in Innate Resistance to Toxoplasma gondii. Cell Host Microbe. 2011;10: 224–236. doi:10.1016/j.chom.2011.07.009

39. Saeij JPJ, Coller S, Boyle JP, Jerome ME, White MW, Boothroyd JC. Toxoplasma co-opts host gene expression by injection of a polymorphic kinase homologue. Nature. 2007;445: 324–327. doi:10.1038/nature05395

40. Wilson HM. SOCS Proteins in Macrophage Polarization and Function. Front Immunol. 2014;5: 357. doi:10.3389/fimmu.2014.00357

41. Wang L, Lu Y-F, Wang C-S, Xie Y-X, Zhao Y-Q, Qian Y-C, et al. HB-EGF Activates the EGFR/HIF-1α Pathway to Induce Proliferation of Arsenic-Transformed Cells and Tumor Growth. Front Oncol. 2020;10. doi:10.3389/fonc.2020.01019

42. Yang ZF, Poon RT, Luo Y, Cheung CK, Ho DW, Lo CM, et al. Up-Regulation of Vascular Endothelial Growth Factor (VEGF) in Small-for-Size Liver Grafts Enhances Macrophage Activities through VEGF Receptor 2-Dependent Pathway1. J Immunol. 2004;173: 2507– 2515. doi:10.4049/jimmunol.173.4.2507

43. Cheng L, Rahman SU, Gong H-Y, Mi R-S, Huang Y, Zhang Y, et al. Transcriptome analysis of a newly established mouse model of Toxoplasma gondii pneumonia. Parasit Vectors. 2023;16: 59. doi:10.1186/s13071-022-05639-3

44. Wood G, Johnson R, Powell J, Bryant OJ, Lastovka F, Brember M, et al. Salmonella injectisome penetration of macrophages triggers rapid translation of transcription factors and protection against cell death. bioRxiv; 2024. p. 2023.07.21.550113. doi:10.1101/2023.07.21.550113

45. Gaji RY, Behnke MS, Lehmann MM, White MW, Carruthers VB. Cell cycle-dependent, intercellular transmission of Toxoplasma gondii is accompanied by marked changes in parasite gene expression. Mol Microbiol. 2011;79: 192–204. doi:10.1111/j.1365-2958.2010.07441.x

46. 46. Cell Counter. [cited 1 Sep 2021]. Available: https://imagej.nih.gov/ij/plugins/cell-counter.html

47. Livak KJ, Schmittgen TD. Analysis of relative gene expression data using real-time quantitative PCR and the 2(-Delta Delta C(T)) Method. Methods San Diego Calif. 2001;25: 402–408. doi:10.1006/meth.2001.1262

48. Smith TS, Heger A, Sudbery I. UMI-tools: Modelling sequencing errors in Unique Molecular Identifiers to improve quantification accuracy. Genome Res. 2017; gr.209601.116. doi:10.1101/gr.209601.116

49. BBMap. In: SourceForge [Internet]. [cited 22 Sep 2021]. Available: https://sourceforge.net/projects/bbmap/

50. Dobin A, Davis CA, Schlesinger F, Drenkow J, Zaleski C, Jha S, et al. STAR: ultrafast universal RNA-seq aligner. Bioinformatics. 2013;29: 15–21. doi:10.1093/bioinformatics/bts635

51. Anders S, Pyl PT, Huber W. HTSeq—a Python framework to work with high-throughput sequencing data. Bioinformatics. 2015;31: 166–169. doi:10.1093/bioinformatics/btu638

52. Love MI, Huber W, Anders S. Moderated estimation of fold change and dispersion for RNA-seq data with DESeq2. Genome Biol. 2014;15: 550. doi:10.1186/s13059-014-0550-8

53. Gaujoux R, Seoighe C. A flexible R package for nonnegative matrix factorization. BMC Bioinformatics. 2010;11: 367. doi:10.1186/1471-2105-11-367

54. Kolberg L, Raudvere U, Kuzmin I, Vilo J, Peterson H. gprofiler2 -- an R package for gene list functional enrichment analysis and namespace conversion toolset g:Profiler. F1000Research. 2020;9: ELIXIR-709. doi:10.12688/f1000research.24956.2

55. Gajria B, Bahl A, Brestelli J, Dommer J, Fischer S, Gao X, et al. ToxoDB: an integrated Toxoplasma gondii database resource. Nucleic Acids Res. 2008;36: D553–D556. doi:10.1093/nar/gkm981

56. Bioicons - high quality science illustrations. [cited 18 Jan 2025]. Available: https://bioicons.com/

