## Supplementary for "*Toxoplasma* strikes preemptively to swiftly suppress macrophage immune response during active infection"

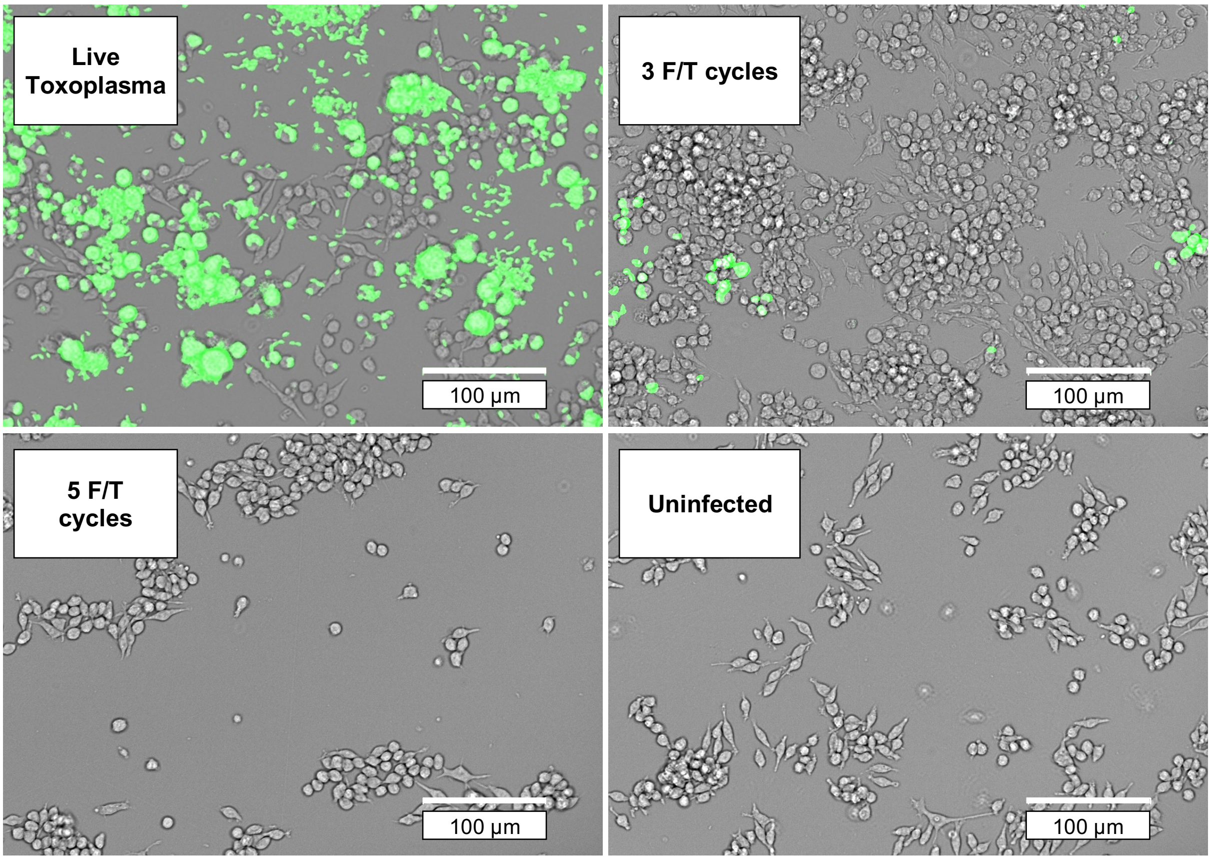


**Figure S1.** Validation of *Toxoplasma gondii* inactivation by freeze-thawing. Green fluorescence overlay indicates *T. gondii* grown overnight. 5 F/T cycles were sufficient to fully inactivate the tachyzoite preparations.

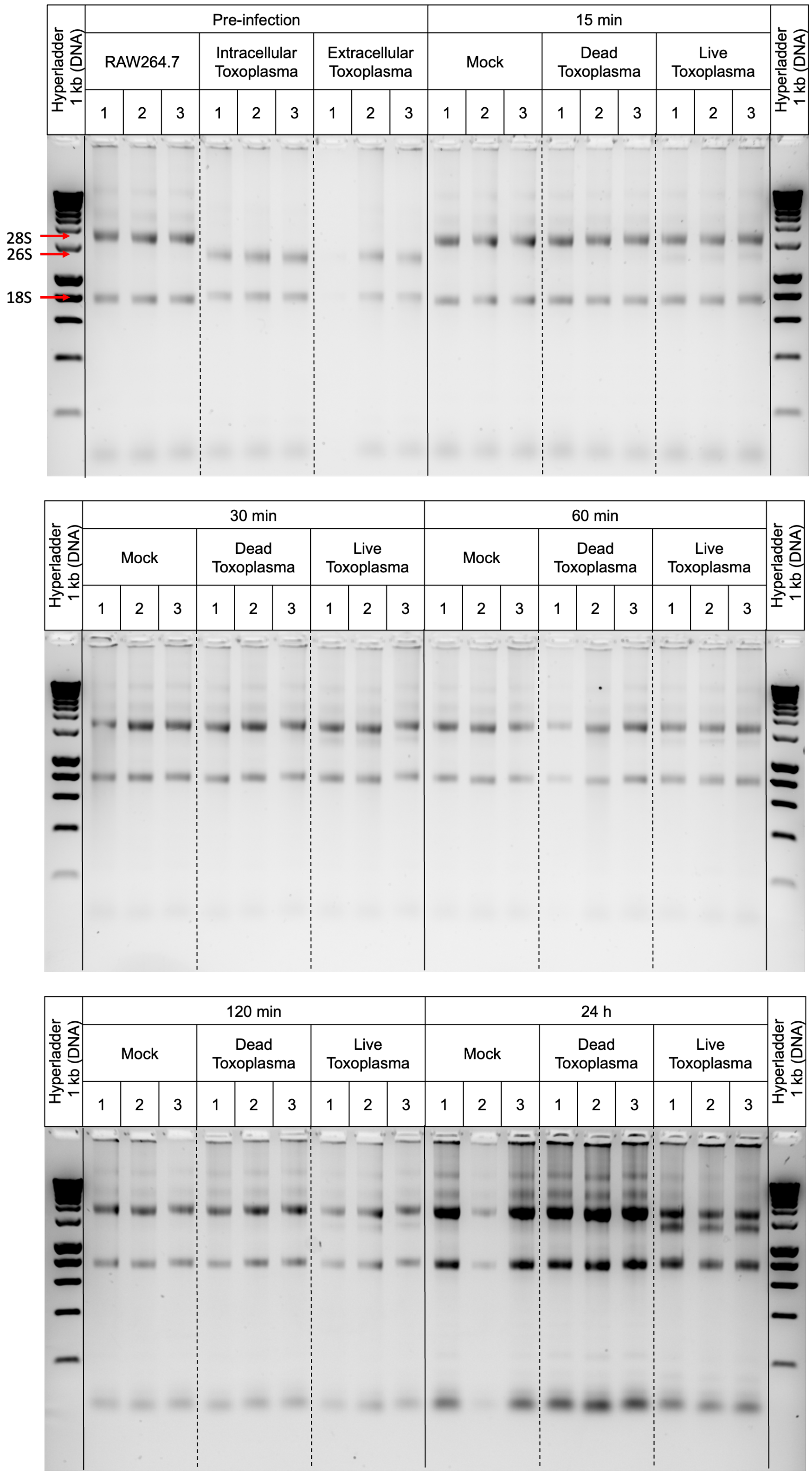


**Figure S2.** Quality control of extracted RNA from time course samples on 2% agarose gels. Expected 28S (mouse), 26S (*T. gondii*) and 18S rRNA band positions are marked with red arrows. By 24 h, 26S rRNA relative amount has increased substantially indicating parasite expansion. Intact rRNA bands and absence of a smear caused by short RNAs indicated sufficient RNA quality.

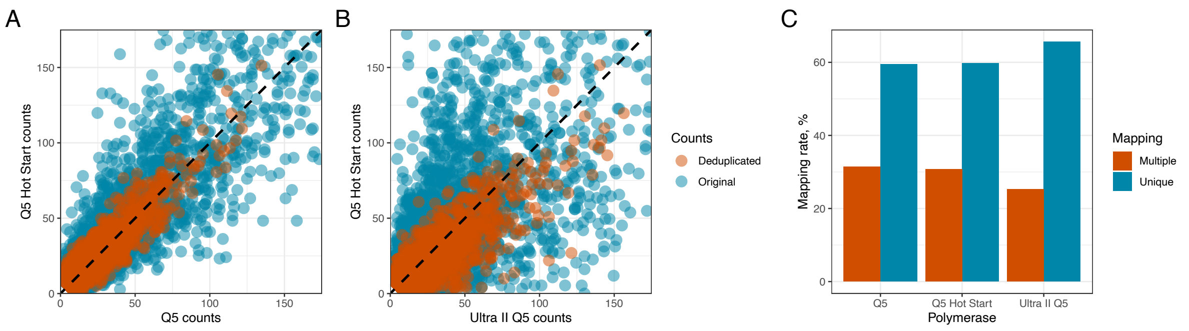


**Figure S3.** Comparison of PARFA-seq library preparation using different PCR reagents. **A** and **B**: Raw and deduplicated count distribution for Q5-Q5 Hot Start and Ultra II Q5-Q5 Hot Start pairs, respectively. **C**: mapping statistics for each library.

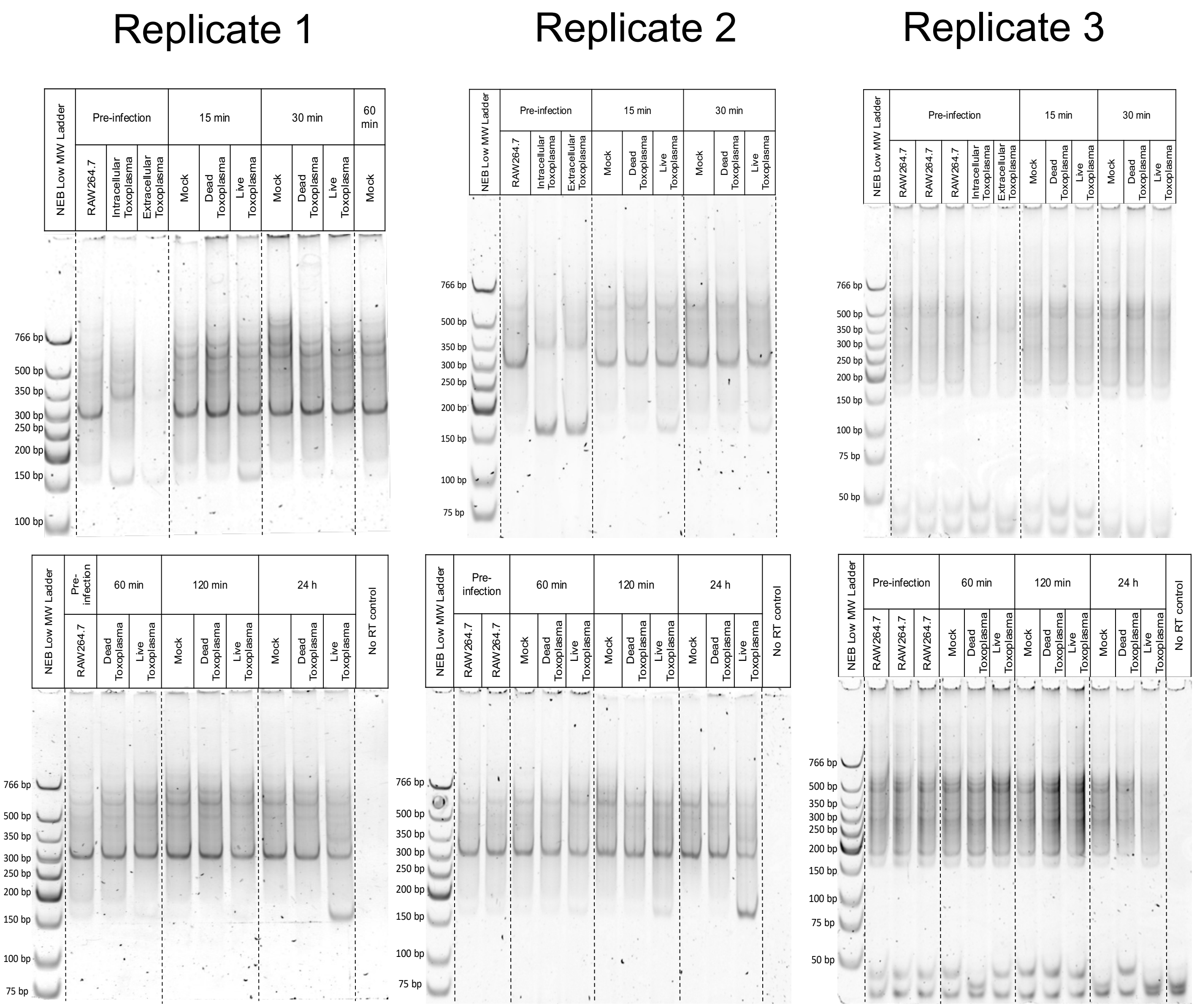


**Figure S4.** PAGE quality control of time course PARFA-seq libraries in preparation before pooling and final size selection.

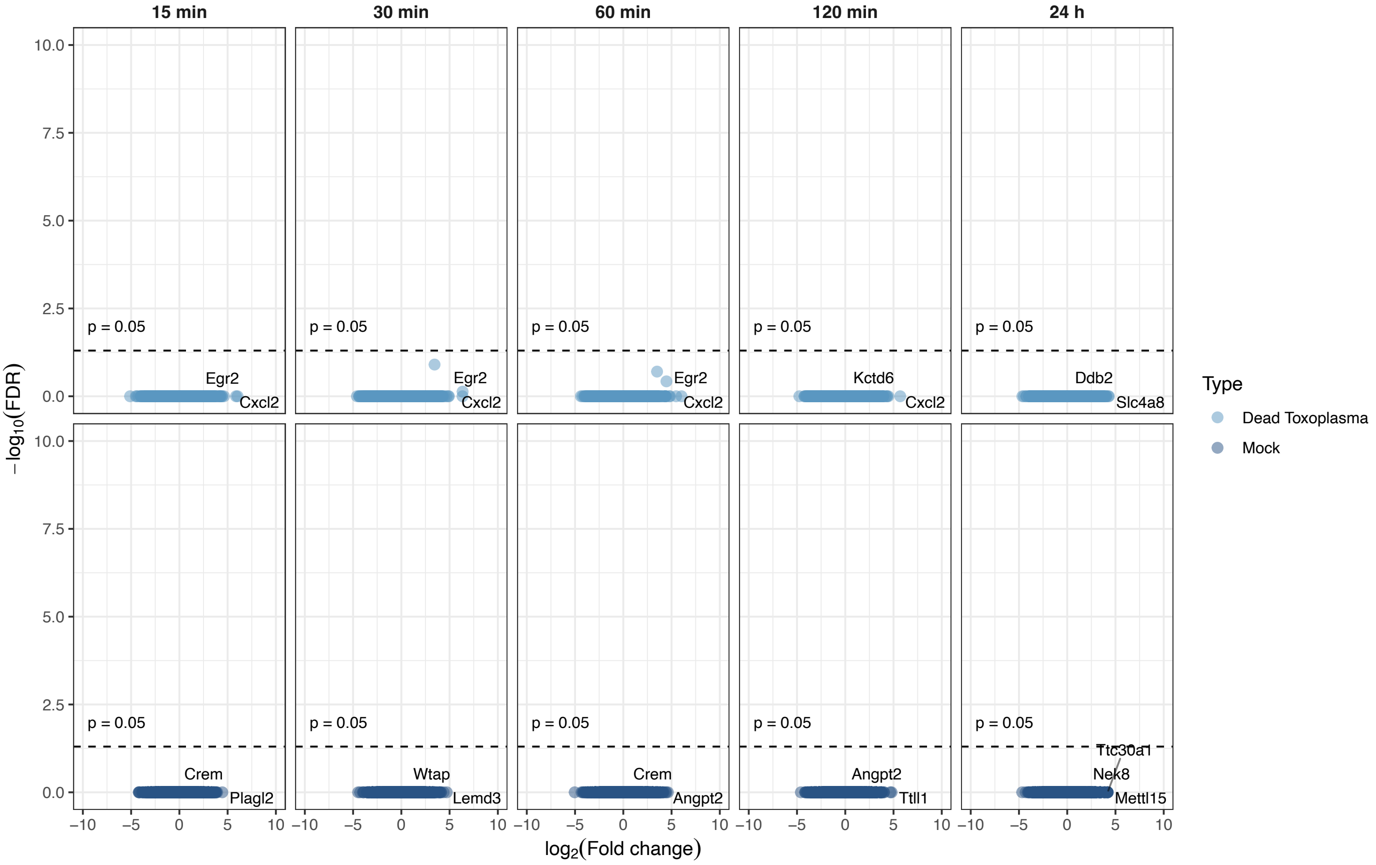


**Figure S5.** Differential expression of Dead Toxoplasma and Mock samples throughout the time course. Dashed line indicates p-value threshold of 0.05. Top upregulated genes are labelled.

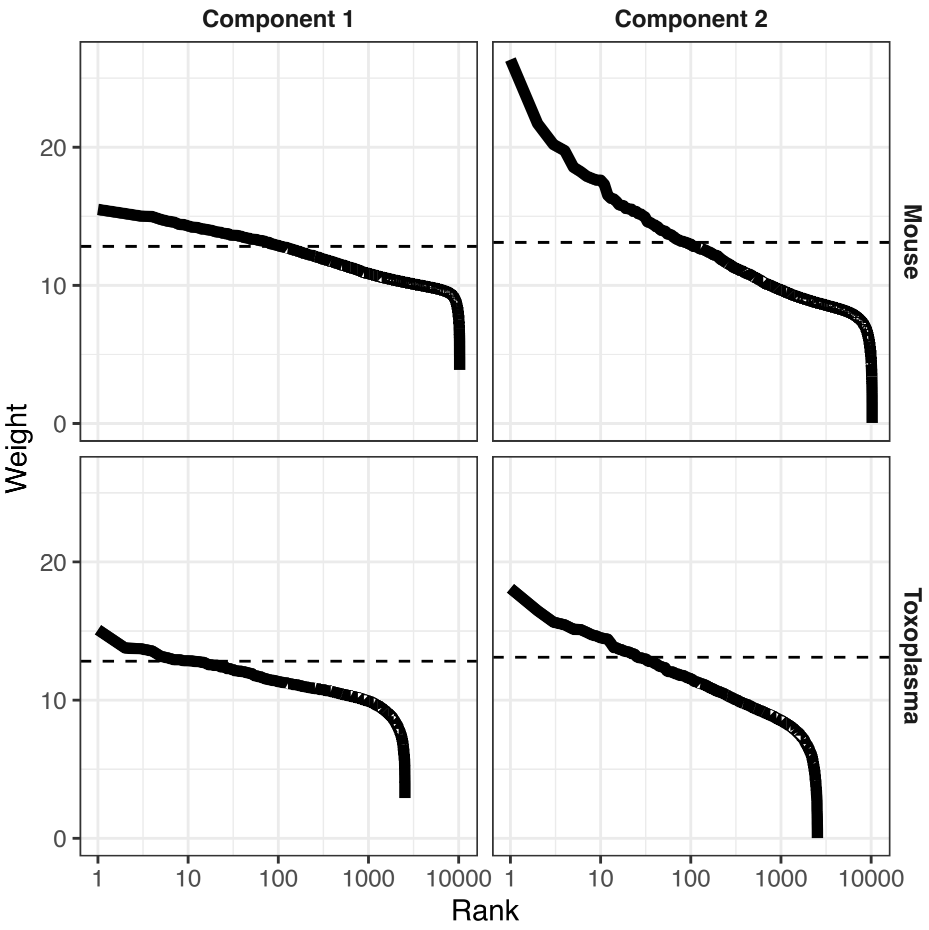


**Figure S6.** Mouse and *T. gondii* genes ranked by weight for each component from non-negative matrix factorization (mouse + *T. gondii* co-transcriptome, 15 min-120 min). Dashed line indicates weight threshold to subset top features for gene set enrichment analysis.

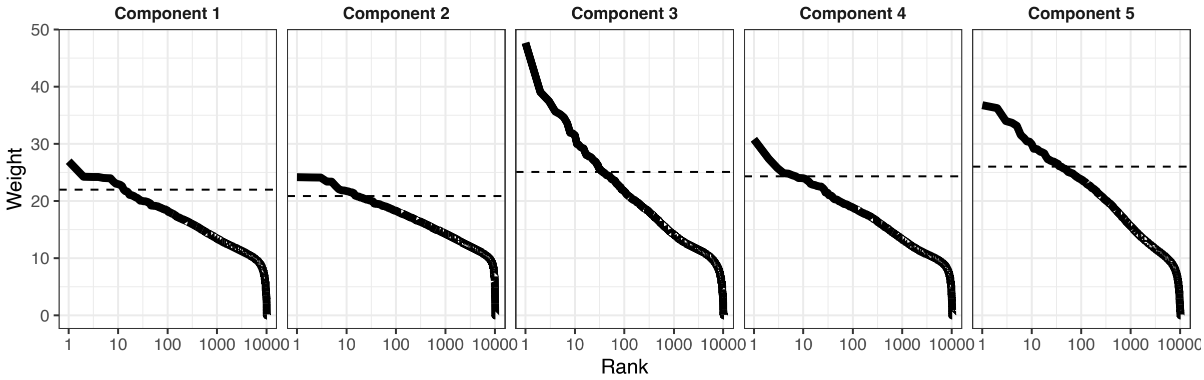


**Figure S7.** Mouse genes ranked by weight for each component from non-negative matrix factorization (mouse transcriptome only, 15 min-24 h). Dashed line indicates weight threshold to subset top features for gene set enrichment analysis.
